# The Value of Zero-filling in In Vivo MRS

**DOI:** 10.1101/2022.05.31.494144

**Authors:** Saipavitra Murali-Manohar, Georg Oeltzschner, Peter B. Barker, Richard A.E. Edden

## Abstract

Two opinions currently exist on the role of zero-filling in data processing for in vivo MRS: that it results in a purely cosmetic interpolation; or that it confers a benefit. Most commonly, in vivo MRS data are acquired as complex time-domain half-echoes, that are Fourier transformed to the give a real spectrum that is modeled for quantification. In this manuscript, we highlight that performing zero-filling draws the independent information from the imaginary part of the spectrum into the real spectrum, improving modeling accuracy. In order to demonstrate this, 10,000 time-domain datasets were simulated as decaying exponentials and noise was added. Data were then Fourier transformed with no-zero filling, 2x, 4x and 8x zero-filling. All spectra were then modeled using a simple single-Lorentzian frequency-domain model. It was demonstrated that 2x zero-filling results in a ∼√2 benefit in modeling accuracy, compared to no zero-filling. There was no additional advantage for further zero-filling.

## Dear Editor

We are writing to highlight the value of zero-filling in *in vivo* MRS. Zero-filling is the practice of appending zeroes to the end of the free induction decay signal prior to Fourier transformation (typically doubling the number of data points, but sometimes more) [1]. Traditionally this has been done to improve the digital resolution of the frequency domain spectrum, and is particularly useful when pulse sequence requirements (e.g. short-TR MR spectroscopic imaging, as one example) limit the length of the data acquisition window.

Zero-filling has been performed from the early days of FT NMR spectroscopy. More recently, it is not widely appreciated within the MRS community (based on discussions with MRS colleagues at ISMRM 2022) what zero-filling achieves, and we believe that it is important that this foundational information is not lost from the *in vivo* MRS community. *There are currently two contradictory opinions on zero-filling:* either that it is a purely cosmetic operation equivalent to interpolation, simply aiding visualization; or that it provides some benefit to the information content and therefore the quantification of MR spectra. In this letter, we show that the latter is correct.

This discussion is limited to the commonest case for in vivo MRS, in which: a complex time-domain half-echo is acquired in quadrature; and real spectra are modeled as a linear combination of frequency-domain basis functions after (possible zero-filling,) Fourier transformation and phasing. In order to demonstrate the benefit of zero-filling, 10,000 time-domain datasets were simulated as decaying complex exponentials. For each simulated acquisition, 1024 datapoints sampled at 1 kHz were simulated with a signal amplitude varied randomly between 0.8 and 1.2 (uniformly distributed), an offset frequency varied randomly between -500 and 500 Hz (uniformly distributed) and a T_2_* decay constant of 100 ms. Noise with a relative amplitude of 30% was added in the time domain, with real and imaginary noise sampled independently from a Gaussian distribution.

Data were Fourier-transformed either to 1024 complex frequency-domain points (no zero-filling) or to 2048, 4096 or 8192 complex frequency-domain points (2x, 4x and 8x zero-filling). The resulting spectra were than modeled using a simple single-Lorentzian frequency-domain model, with three parameters corresponding to amplitude, offset frequency and width.

Figure 1A shows the real part of one of the spectra, overlaid by the Lorentzian model, illustrating the SNR of 3.67. Figure 1B shows the results of the simulations, plotting the modeled amplitude against the correct amplitude for the zero-filled and non zero-filled cases. It can be clearly seen by eye that the zero-filled modeling performs better, with a tighter distribution about the diagonal. The standard deviation of the modeling errors were 6.2% in the zero-filled case and 8.9% in the non-zero-filled case, equivalent to a ∼√2 improvement. This benefit is maintained, but not increased by further zero-filling, as shown in Figure 1C.

**Figure 1:**
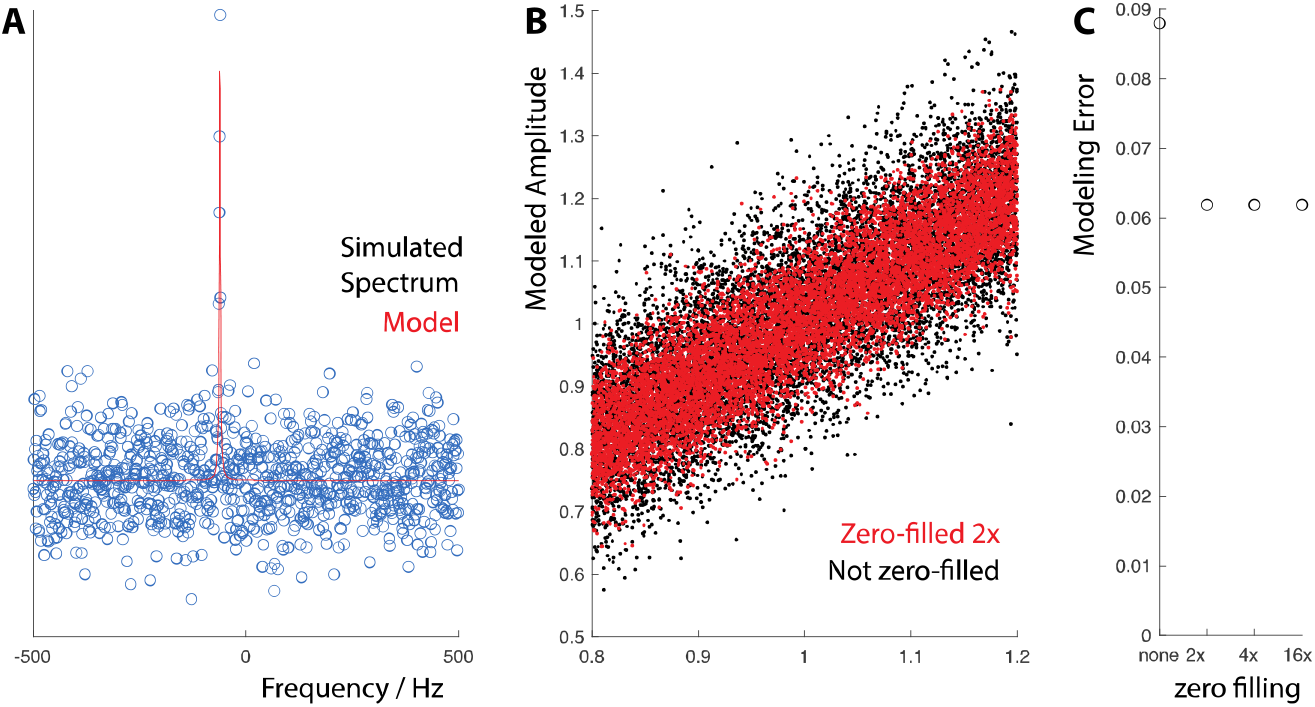
A) Simulated spectrum with noise, modeled by a real Lorentzian model. B) Results of 10,000 simulated models, clearly showing the reduced errors seen when modeling zero-filled spectra. C) Plot of modeling error against zero-filling factor, indicating a 1/√2 reduction from the first zero-fill and no further benefit thereafter.

If zero-filling cannot add information to the data, how can it improve the accuracy of modeling? The real and imaginary spectrum that arise from a complex Fourier transform of half-echo data are not Hilbert pairs (i.e. they contain independent information and are not mutually predictable by the Hilbert transform) *until zero-filling has been performed*, as originally emphasized for NMR [1]. Before zero-filling the information content of the real and imaginary spectra is distinct, and after zero-filling, it is not. The first zero-fill thus has the effect of drawing the independent information stored in the imaginary spectrum into the real spectrum. Although it appears to result in interpolation, the information is not predictable from the real spectrum, but rather added from the imaginary spectrum. The benefit in terms of modeling accuracy is that twice as many independent points sample each signal of interest, conferring a ∼√2 benefit in integrated SNR [3].

In closing, we highlight an important benefit of zero-filling in MRS applications where only the real spectrum is to be analyzed. Zero-filling is purely cosmetic after the first interpolation, but the first 2x zero-fill confers a ∼√2 benefit in modeling accuracy.

## Supplementary material

Full Matlab code to repeat the calculation performed and produce Figure 1

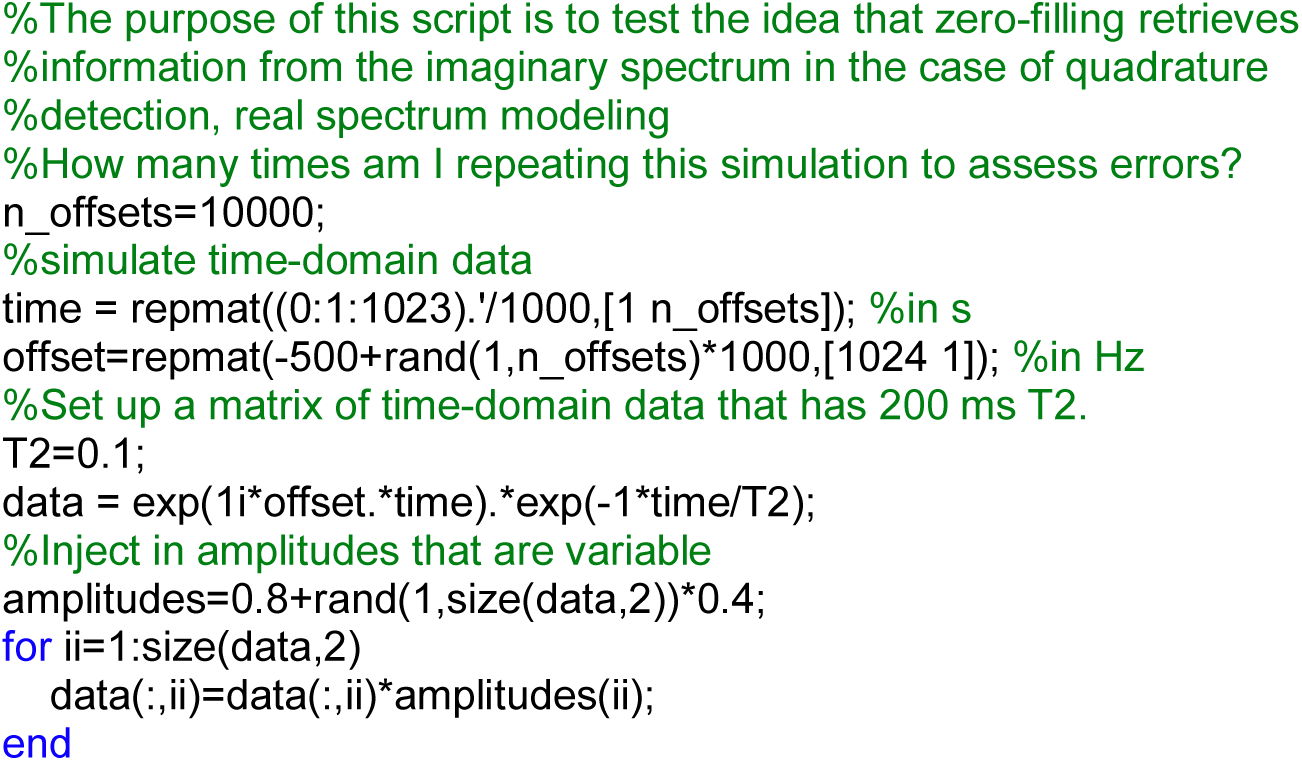
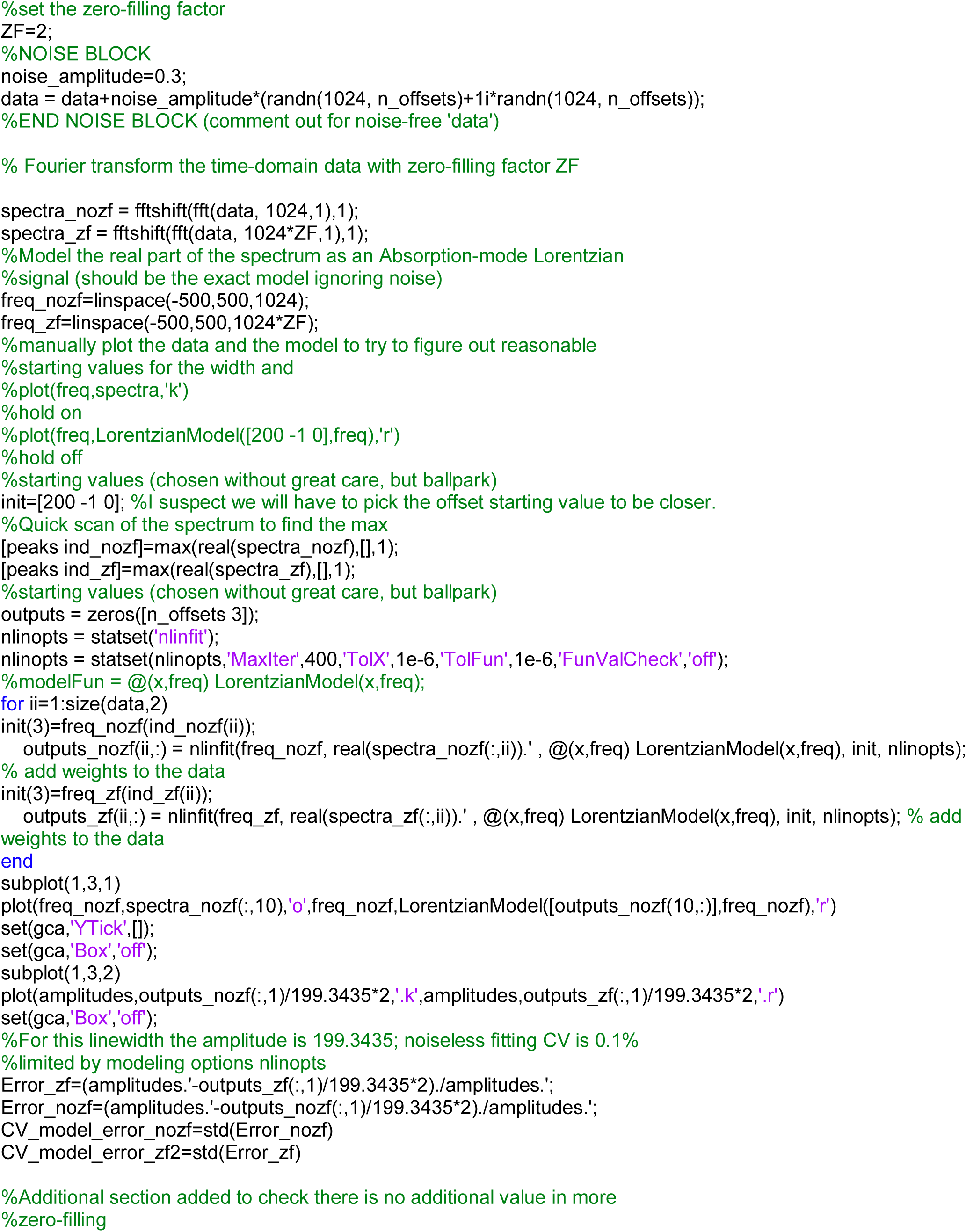
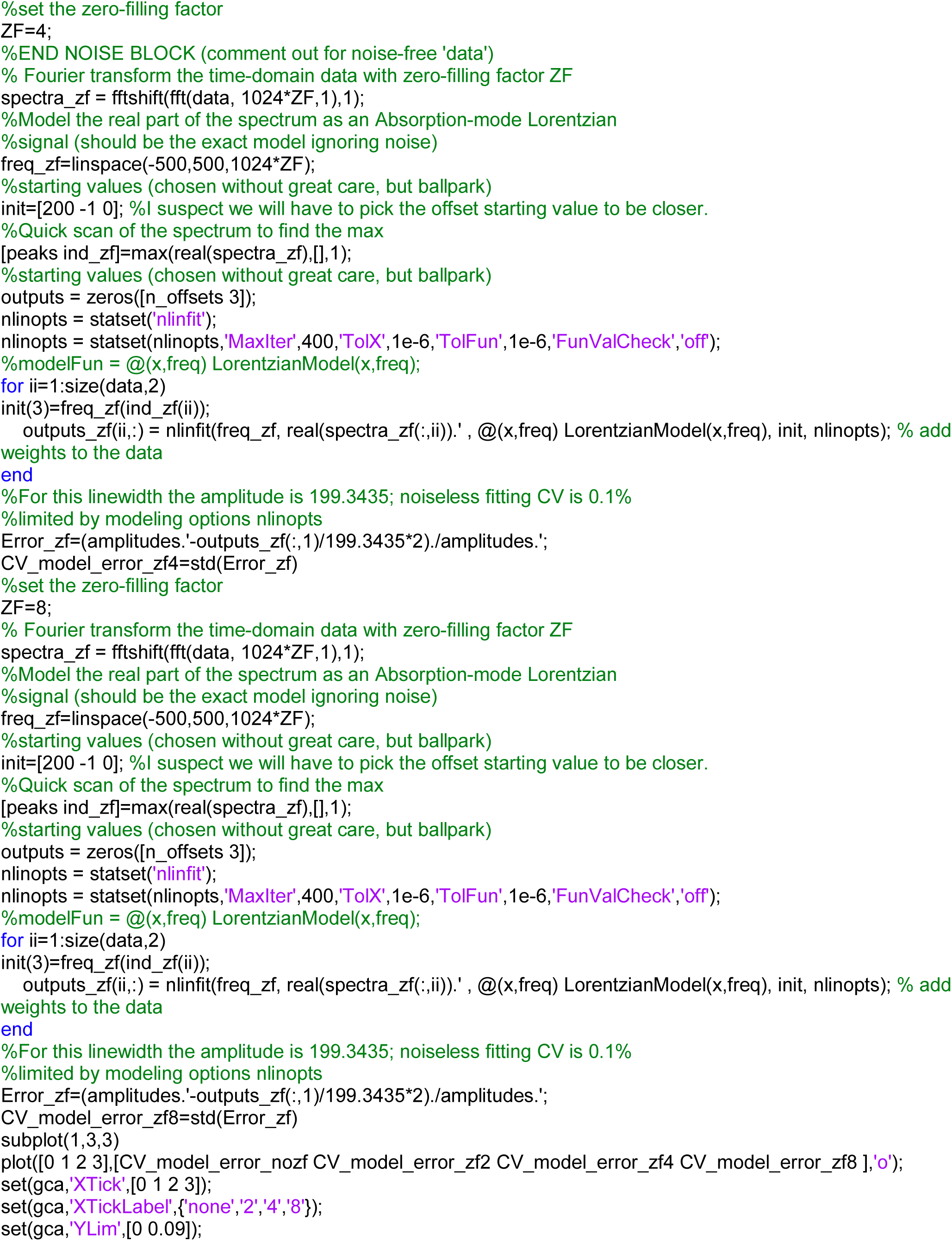
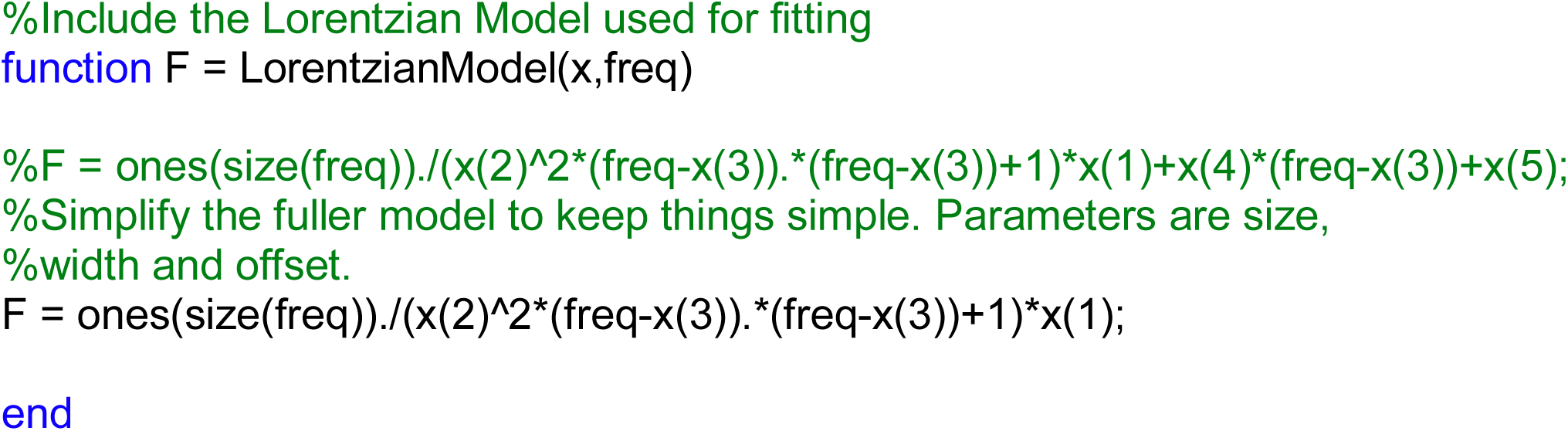

